# Triphosphates of the Two Components in DESCOVY and TRUVADA are Inhibitors of the SARS-CoV-2 Polymerase

**DOI:** 10.1101/2020.04.03.022939

**Authors:** Steffen Jockusch, Chuanjuan Tao, Xiaoxu Li, Thomas K. Anderson, Minchen Chien, Shiv Kumar, James J. Russo, Robert N. Kirchdoerfer, Jingyue Ju

**Affiliations:** Center for Genome Technology and Biomolecular Engineering, Columbia University, New York, NY 10027; Departments of Chemistry, Columbia University, New York, NY 10027; Departments of Chemical Engineering, Columbia University, New York, NY 10027; Departments of Pharmacology, Columbia University, New York, NY 10027; Departments of Biochemistry, University of Wisconsin-Madison, Madison, WI 53706; Institute of Molecular Virology, University of Wisconsin-Madison, Madison, WI 53706

## Abstract

SARS-CoV-2, a member of the coronavirus family, is responsible for the current COVID-19 pandemic. We previously demonstrated that four nucleotide analogues (specifically, the active triphosphate forms of Sofosbuvir, Alovudine, AZT and Tenofovir alafenamide) inhibit the SARS-CoV-2 RNA-dependent RNA polymerase (RdRp). Tenofovir and emtricitabine are the two components in DESCOVY and TRUVADA, the two FDA-approved medications for use as pre-exposure prophylaxis (PrEP) to prevent HIV infection. This is a preventative method in which individuals who are HIV negative (but at high-risk of contracting the virus) take the combination drug daily to reduce the chance of becoming infected with HIV. PrEP can stop HIV from replicating and spreading throughout the body. We report here that the triphosphates of tenofovir and emtricitabine, the two components in DESCOVY and TRUVADA, act as terminators for the SARS-CoV-2 RdRp catalyzed reaction. These results provide a molecular basis to evaluate the potential of DESCOVY and TRUVADA as PrEP for COVID-19.

## Introduction

The COVID-19 pandemic, caused by the novel coronavirus SARS-CoV-2, has now reached critical levels worldwide and is still spreading. SARS-CoV-2 is a member of the subgenus *Sarbecovirus* in the Orthocoronavirinae subfamily.^1^ Coronaviruses are single-strand RNA viruses that share properties with other RNA viruses such as the hepatitis C virus (HCV), West Nile virus, Marburg virus, HIV, Ebola virus, dengue virus, and rhinoviruses. Coronaviruses, HCV and the flaviviruses are positive-sense single-strand RNA viruses^2,3^ that share a similar replication mechanism requiring an RNA-dependent RNA polymerase (RdRp) catalyzed reaction.

Although potential drug candidates have been designed to target nearly every stage of the viral infection,^2^ no effective drug is currently approved to treat COVID-19. RdRp in coronaviruses is a precise and well-defined drug target; the active site of the RdRp is highly conserved among different positive-sense RNA viruses and shares common structural features in these viruses.^4^ RdRps, including the coronavirus polymerase, have low fidelity,^5^ which enables them to recognize a variety of modified nucleotide analogues as substrates. Such nucleotide and nucleoside analogues may inhibit further RNA-polymerase catalyzed RNA replication and are therefore important candidate anti-viral agents.^6-9^

In our previous studies, we used molecular analysis to design and select nucleotide analogues as potential inhibitors of the SARS-CoV-2 RdRp,^10,11^ with particular focus on two key properties: (1) their similarity in size and structure to natural nucleotides, including their ability to bind within the active site of the polymerase and (2) their ability to terminate polymerase reactions due either to (a) the presence of blocking groups on the 3’-OH or (b) the presence of chemical modification groups at nearby positions on the sugar ring. Based on these criteria, we previously tested the active triphosphate forms of Sofosbuvir, Alovudine and AZT with the RdRps of both SARS-CoV and SARS-CoV-2. These three triphosphates, 2’-F,Me-UTP, 3’-F-dTTP and 3’-N3-dTTP, all demonstrated the ability to be incorporated by these two coronavirus RdRps, and to block further incorporation.^10,11^ We also demonstrated the ability of the active triphosphate form of tenofovir, tenofovir diphosphate (TFV-DP), to inhibit the SARS-CoV-2 RdRp.^11^ Tenofovir prodrugs are often used in combination with the drug emtricitabine as anti-viral medications. We report here that the active triphosphate forms of both tenofovir and emtricitabine are inhibitors of the SARS-CoV-2 RdRp catalyzed reaction in side-by-side experiments.

Tenofovir diphosphate (TFV-DP) and Emtricitabine triphosphate (Ec-TP) (Fig. 1 c, g) are the active triphosphate forms of two viral reverse transcriptase inhibitors for HIV and hepatitis B virus (HBV). Gilead’s TRUVADA and DESCOVY are two FDA-approved medications for use as pre-exposure prophylaxis (PrEP) to prevent HIV infection. PrEP is a preventative method in which individuals who are HIV negative (but at high-risk of contracting the virus) take the combination drug daily to reduce the chance of becoming infected with HIV.^12-15^ Both TRUVADA and DESCOVY consist of the prodrugs, Emtricitabine (2’,3’-dideoxy-5-fluoro-3’-thiacytidine, FTC) and either of two prodrug forms of Tenofovir (Tenofovir disoproxil fumarate (TDF) in TRUVADA and Tenofovir alafenamide (TAF) in DESCOVY). Overall, the combination drugs are well tolerated, with a small but significant reduction in the number of patients with side effects in the latter.^16^ These drugs are often included in the cocktails for patients with HIV/AIDS. It is critical to ensure that those being administered these drugs for PrEP are HIV negative. Resistance mutations have been observed in a small percentage of individuals using FTC plus TDF.^17^ It is therefore important to screen them for seroconversion and possible drug resistance.

**Fig. 1.**
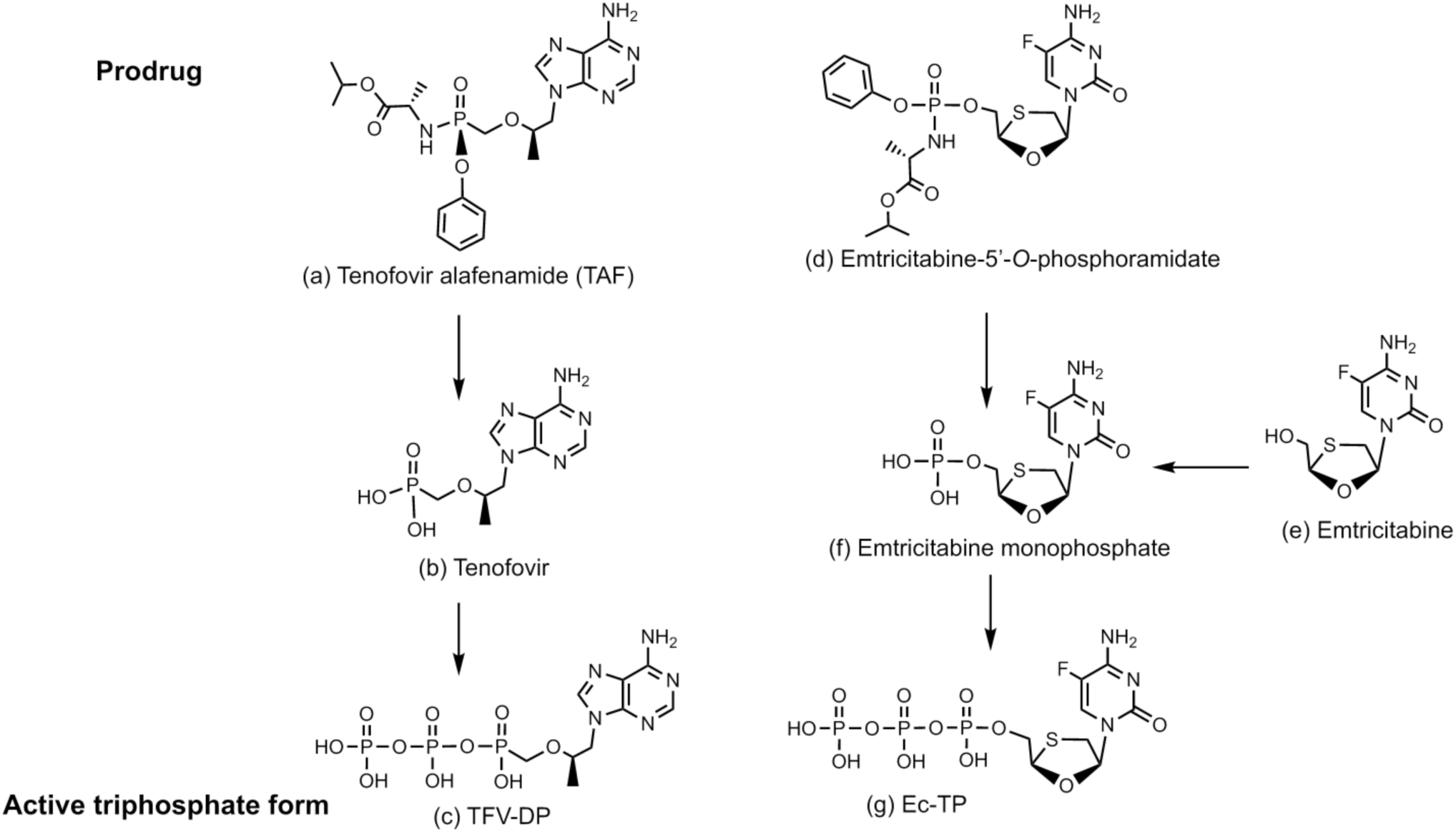
Structures of prodrug viral inhibitors. Prodrugs Tenofovir alafenamide (TAF) (a), Emtricitabine-5’-O-phosphoramidate (d) and Emtricitabine (e), their monophosphate forms Tenofovir (TFV) and Emtricitabine monophosphate (b and f, respectively), and their active triphosphate forms (c and g, respectively).

Tenofovir alafenamide (TAF, Fig. 1a), a prodrug form of TFV-DP, is activated by a series of hydrolases to the deprotected monophosphate form, Tenofovir (TFV) (Fig. 1b), and then by two consecutive kinase reactions to TFV-DP (Fig. 1c).^18^ TAF shows potent activity for HIV and HBV, but only limited inhibition of host nuclear and mitochondrial polymerases.^19,20^ TFV-DP is an acyclic nucleotide and notably does not have a 3’-OH group. Upon incorporation by both HIV and HBV polymerase catalyzed reactions, nucleic acid elongation is terminated and viral replication is stopped.^18,19^ In addition, resistance mutations rarely occur in individuals treated with regimens that include TAF.^21^ TDF is another prodrug of TFV-DP, differing only in the protective group used to mask the phosphate in comparison to TAF^18^. Given that Tenofovir diphosphate (TFV-DP), which is the common active triphosphate form of TAF or TDF, is much smaller than natural nucleoside triphosphates, we expect that it can easily fit within the active site of SARS-CoV-2 RdRp. As an acyclic nucleotide, TFV-DP lacks a normal sugar ring configuration, thus we reasoned that it is unlikely to be recognized by 3’-exonucleases involved in SARS-CoV-2 proofreading processes, thereby decreasing the likelihood of developing resistance to inhibition by the drug. Based on these analyses, we previously demonstrated the the ability of TFV-DP to inhibit the SARS-CoV-2 RdRp.^11^ This inhibitory effect was furthered confirmed in the current study.

Emtricitabine (FTC) is a 5-fluorocytidine analog containing an oxathiolane ring with an unnatural (-)-β-*L*-stereochemical configuration (Fig. 1e).^22^ This pro-drug is converted by cellular enzymes, first to a monophosphate (Fig. 1f) then to the active triphosphate form Ec-TP (Fig. 1g). An alternative 5’-*O*-phosphoramidate prodrug can also be synthesized (Fig. 1d). The absence of an OH group at the 3’ position ensures that once this nucleotide analogue is incorporated into the primer in polymerase reaction, no further incorporation of nucleotides by the polymerase can occur. Like TAF, FTC is effective against the retrovirus HIV and has also been shown to inhibit HBV in clinical studies.^23^ Due to its structural similarity with molecules we previously tested^11^ and because FTC is a key component of DESCOVY and TRUVADA, Ec-TP (the active form of FTC) was included in this SARS-CoV-2 RdRp inhibitory study.

## Results and Discussion

We tested the ability of the active triphosphate forms of both tenofovir and emtricitabine to be incorporated by the SARS-CoV-2 RdRp. The RdRp of SARS-CoV-2 (referred to as nsp12) and its two protein cofactors (nsp7 and nsp8), whose homologs were shown to be required for the processive polymerase activity of nsp12 in SARS-CoV,^24,25^ were cloned and purified as previously described.^11^ We then performed polymerase extension assays with UTP + TFV-DP, and UTP + ATP + Ec-TP, following the addition of a pre-annealed RNA template and primer to a pre-assembled mixture of the SARS-CoV-2 RdRp (nsp12) and the two cofactor proteins (nsp7 and nsp8). The primer extension products from the reaction were analyzed by MALDI-TOF-MS. The RNA template and primer, corresponding to the 3’ end of the SARS-CoV-2 genome, were used for the polymerase assay; their sequences are indicated at the top of Fig. 2. In the case of tenofovir diphosphate (TFV-DP) that contains an adenine base, because there are two A’s in a row followed by a U in the next available positions of the template for RNA polymerase extension downstream of the priming site, we added both UTP and TFV-DP in the extension reaction. Based on results we previously obtained for TFV-DP,^11^ we reduced the ratio of UTP and TFV-DP from 1:10 to 1:100 in this current experiment. If TFV-DP acts as a permanent terminator of the RdRp reaction, we would ideally observe a peak in the spectrum at the third primer extension position caused by incorporation of two Us and one TFV-DP. In the case of emtricitabine triphosphate (Ec-TP), which is a C analogue, we need to extend the primer to the first available G in the template sequence. Since the next available positions in the template consist of two As, four Us, another A and a G, we included UTP, ATP and Ec-TP in the extension reaction. Thus, if Ec-TP serves as a permanent inhibitor of the RdRp reaction, we would ideally observe a peak in the spectrum at the eighth primer extension position caused by incorporation of two Us, four As, another U and one Ec-TP.

**Fig. 2.**
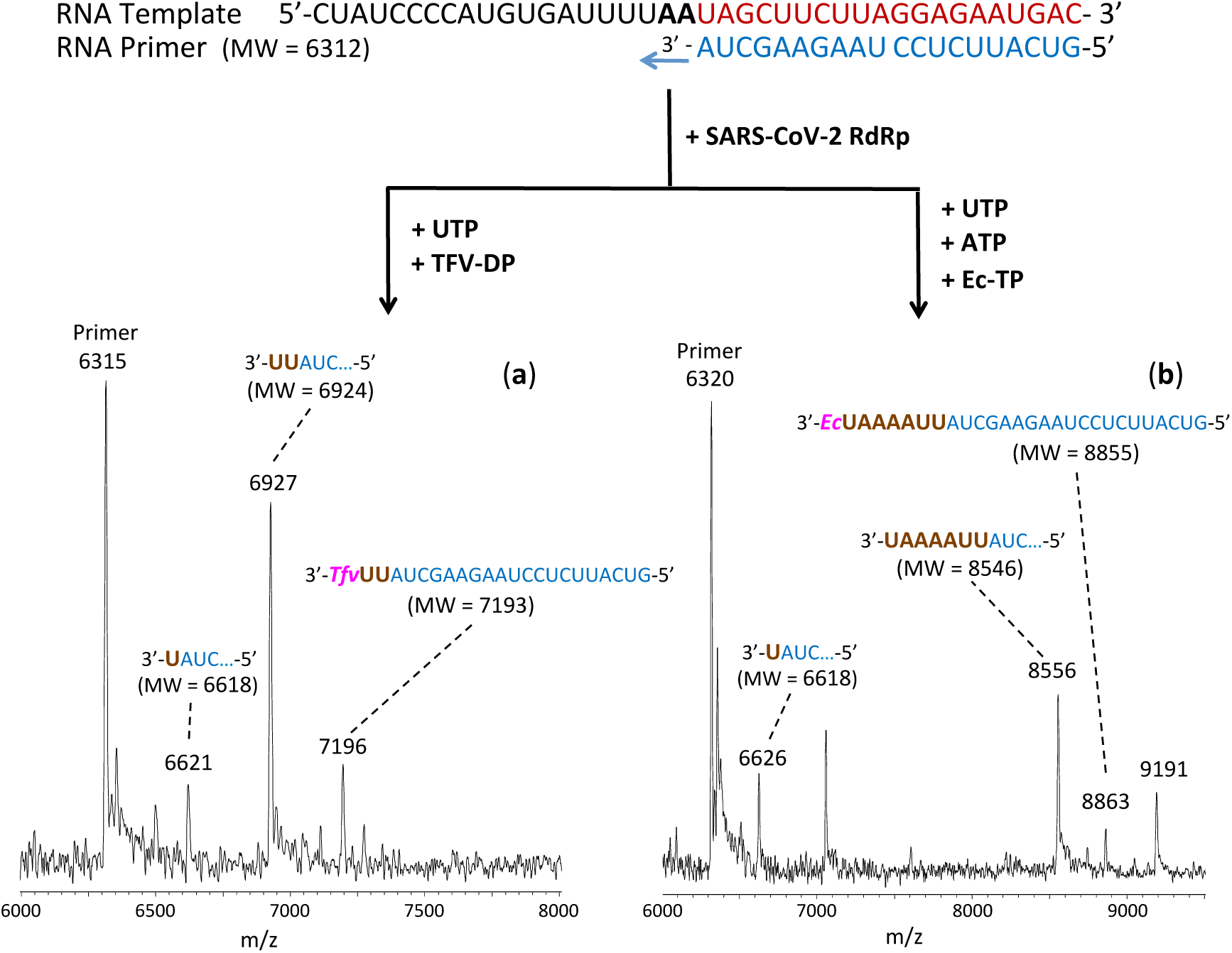
Incorporation of TFV-DP and Ec-TP by SARS-CoV-2 RdRp to terminate the polymerase reaction. The sequences of the primer and template used for these extension reactions, which are at the 3’ end of the SARS-CoV-2 genome, are shown at the top of the figure. Polymerase extension reactions were performed by incubating (**a**) UTP + TFV-DP, and (**b**) UTP + ATP + Ec-TP with pre-assembled SARS-CoV-2 polymerase (nsp12, nsp7 and nsp8), the indicated RNA template and primer, and the appropriate reaction buffer, followed by the detection of reaction products by MALDI-TOF MS. The detailed procedure is described in the Methods section. The accuracy for m/z determination is ± 10 Da.

The results of the MALDI-TOF MS analysis of the primer extension products are shown in Fig. 2. The observed peaks generally fit the predictions above; however, additional peaks from products assigned to intermediate stages of the extension reaction were also observed. In the case of TFV-DP (Fig. 2a), a peak with the expected molecular weight representing incorporation of two Us and one TFV-DP was obtained (7196 Da observed, 7193 Da expected). As anticipated, partial extension products were also observed for a single U (6621 Da observed, 6618 Da expected) and for two Us (6927 Da observed, 6924 Da expected). Most importantly, once the TFV-DP was incorporated, there was no further extension, which confirms that it is a permanent terminator. These results are similar to our previous primer extension reactions with UTP and TFV-DP.^11^ However, we previously observed significant amounts of misincorporation products (3 Us) due to the higher concentration of UTP. Under the current conditions, misincorporations were insignificant (Fig. 2a). In the case of Ec-TP (Fig. 2b), an expected peak indicating extension by three Us, four As and one Ec-TP was observed (8863 Da observed, 8855 Da expected), but shorter extension products also occurred, such as the extension by one U (6626 Da observed, 6618 Da expected), or a longer primer extension product containing the sequence Primer–UUAAAAU-3’ but excluding the incorporation of an Ec-TP (8556 Da observed, 8546 Da expected). The extra peak at the right (9191 Da) could be explained by misincorporation events, such as a longer primer extension product containing the sequence Primer– UUAAAAU*U*A-3’ (9181 Da expected) in which a U (italicized) was incorporated instead of a C, followed by the incorporation of an A. The occurrence of misincorporation indicates that SARS-CoV-2 RdRp has low fidelity, a property similar to other RNA viruses. ^5,26^

In summary, these results demonstrate that the nucleotide analogues TFV-DP and Ec-TP are permanent terminators for the SARS-CoV-2 RdRp catalyzed reaction. Their prodrug versions (TAF or TDF and Emtricitabine or Emtricitabine-5’-*O*-phosphoramidate that can be readily synthesized using the ProTide prodrug approach^27^) can be used as lead compounds for COVID-19 therapeutics development. More importantly, since these molecules are contained in the FDA-approved combination drugs TRUVADA and DISCOVY, our results provide a molecular basis to further evaluate them in SARS-CoV-2 virus inhibition and animal models to test their efficacy for the development of potential COVID-19 medications. These current and future studies may also provide potential options for preventing COVID-19 infection in a manner similar to PrEP for HIV prevention.

## Methods

### Extension reactions with RNA-dependent RNA polymerase

Oligonucleotides were purchased from Integrated DNA Technologies. The primer and template (sequences shown in Fig. 2) were annealed by heating to 70°C for 10 min and cooling to room temperature in 1x reaction buffer. The RNA polymerase mixture consisting of 6 *µ*M nsp12 and 18 *µ*M each of cofactors nsp7 and nsp8 (1:3:3 ratio) was incubated for 15 min at room temperature in 1x reaction buffer. Then 5 *µ*l of the annealed template primer solution containing 2 *µ*M template and 1.7 *µ*M primer in 1x reaction buffer was added to 10 *µ*l of the RNA polymerase mixture and incubated for an additional 10 min at room temperature. Finally, 5 *µ*l of a solution containing either 2 mM TFV-DP and 20 *µ*M UTP (a) or 2 mM Ec-TP (Toronto Research Chemicals), 200 *µ*M UTP and 200 *µ*M ATP (b) in 1x reaction buffer was added, and incubation was carried out for 2 hrs at 30°C. The final concentrations of reagents in the 20 *µ*l extension reactions were 3 *µ*M nsp12, 9 *µ*M nsp7, 9 *µ*M nsp8, 425 nM RNA primer, 500 nM RNA template, either 500 *µ*M TFV-DP / 5 *µ*M UTP or 500 *µ*M Ec-TP / 50 *µ*M UTP / 50*µ*M ATP. The 1x reaction buffer contains the following reagents: 10 mM Tris-HCl pH 8, 10 mM KCl, 2 mM MgCl2 and 1 mM β-mercaptoethanol. Following desalting using an Oligo Clean & Concentrator (Zymo Research), the polymerase reaction products were subjected to MALDI-TOF-MS (Bruker ultrafleXtreme) analysis.

## Acknowledgements

This research is supported by Columbia University, a grant from the Jack Ma Foundation to J.J. and a National Institute of Allergy and Infectious Disease grant AI123498 to R.N.K. A patent application on the work described has been filed.

## Author contributions

J.J. and R.N.K. conceived and directed the project; the approaches and assays were designed and conducted by J.J., X.L., S.K., S.J., J.J.R., M.C. and C.T., and SARS-CoV-2 polymerase (nsp12) and associated proteins (nsp 7 and 8) were cloned and purified by T.K.A. and R.N.K. Data were analyzed by all authors. All authors wrote and reviewed the manuscript.

## References

1. Zhu, N. et al., for the China Novel Coronavirus Investigating and Research Team. A novel coronavirus from patients with pneumonia in China, 2019. N Eng J Med 382, 727–733 (2020).

2. Zumla, A., Chan, J. F. W., Azhar, E. I., Hui, D. S. C. & Yuen, K.-Y. Coronaviruses – drug discovery and therapeutic options. Nat Rev | Drug Discovery 15, 327–347 (2016).

3. Dustin, L. B., Bartolini, B., Capobianchi, M. R. & Pistello, M. Hepatitis C virus: life cycle in cells, infection and host response, and analysis of molecular markers influencing the outcome of infection and response to therapy. Clin Microbiol Infect 22, 826–832 (2016).

4. te Velthuis, A. J. W. Common and unique features of viral RNA-dependent polymerases. Cell Mol Life Sci 71, 4403–4420 (2014).

5. Selisko, B., Papageorgiou, N., Ferron, F. & Canard, B. Structural and functional basis of the fidelity of nucleotide selection by *Flavivirus* RNA-dependent RNA polymerases. Viruses 10, 59 (2018).

6. McKenna, C. E. et al. Inhibitors of viral nucleic acid polymerases. Pyrophosphate analogues. ACS Symposium Series 401, 1–16. Chapter 1 (1989).

7. Öberg, B. Rational design of polymerase inhibitors as antiviral drugs. Antiviral Res 71, 90–95 (2006).

8. Eltahla, A. A., Luciani, F., White, P. A., Lloyd, A. R. & Bull, R. A. Inhibitors of the hepatitis C virus polymerase; mode of action and resistance. Viruses 7, 5206–5224 (2015).

9. De Clercq, E. & Li, G. Approved antiviral drugs over the past 50 years. Clin Microbiol Rev 29, 695–747 (2016).

10. Ju, J. et al. Nucleotide analogues as inhibitors of SARS-CoV polymerase. bioRxiv preprint. doi.org/10.1101/2020.03.12.989186 (2020).

11. Chien, M. C. et al. Nucleotide analogues as inhibitors of SARS-CoV-2 polymerase. bioRxiv preprint. doi.org/10.1101/2020.03.18.997585 (2020).

12. Anderson, P. L., Kiser, J. J., Gardner, E. M., Rower, J. E., Meditz, A. & Grant, R. M. Pharmacological considerations for tenofovir and emtricitabine to prevent HIV infection. J Antimicrob Chemother 66, 240–250 (2011).

13. Grant, R. M. et al. Preexposure chemoprophylaxis for HIV prevention in men who have sex with men. N Eng J Med 363, 2587–2599 (2010).

14. Anderson, P. L. et al. Emtricitabine-tenofovir concentrations and pre-exposure prophylaxis efficacy in men who have sex with men. Sci Transl Med 4, 151ra125 (2012).

15. McNairy, M. L. & El-Sadr, W. M. A paradigm shift: focus on the HIV prevention continuum. Clin Infect Dis 59, S12–S15 (2014).

16. Spinner, C. D. et al. DISCOVER study for HIV pre-exposure prophylaxis (PrEP): F/TAF has a more rapid onset and longer sustained duration of HIV protection compared with F/TDF. IAS 2019 Abstract. http://programme.ias2019.org/Abstract/Abstract/4898.

17. Parikh, U. M. & Mellors, J. W. Should we fear resistance from tenofovir/emtricitabine preexposure prophylaxis? Curr Opin HIV AIDS 11, 49–55 (2016).

18. De Clercq, E. Tenofovir alafenamide (TAF) as the successor of tenofovir disoproxil fumarate (TDF). Biochem Pharmacol. doi.org/10.1016/j.bcp.2016.04.015 (2016).

19. Birkus, G. B. Intracellular activation of Tenofovir alafenamide and the effect of viral and host protease inhibitors. Antimicrob Agents Chemother 60, 316–322 (2016).

20. Lou, L. Advances in nucleotide antiviral development from scientific discovery to clinical applications: Tenofovir disproxil fumarate for hepatitis B. J Clin Translat Hepatol 1, 33–38 (2013).

21. Margot, N. et al. Rare emergence of drug resistance in HIV-1 treatment-naïve patients receiving elvitegravir/cobicistat/emtricitabine/tenofovir alafenamide for 144 weeks. J Clin Virol 103, 37–42 (2018).

22. Hung, M., Tokarsky, E. J., Lagpacan L., Zhang, L., Suo, Z. & Lansdon, E. B. Elucidating molecular interactions at *L-*nucleotides wih HIV-1 reverse transcriptase and mechanism of M184V-caused drug resistance. Commun Biol 2, 469 (2019).

23. Lim, S. G. et al. A double-blind placebo-controlled study of emtricitabine in chronic hepatitis B. Arch Intern Med 166, 49–56 (2006).

24. Subissi, L. et al. One severe acute respiratory syndrome coronavirus protein complex integrates processive RNA polymerase and exonuclease activities. Proc Natl Acad Sci USA 111, E3900–E3909 (2014).

25. Kirchdoerfer, R. N. & Ward, A. B. Structure of the SARS-CoV nsp12 polymerase bound to nsp7 and nsp8 co-factors. Nature Commun 10, 2342 (2019).

26. Arnold, J. J., Vignuzzi, M., Stone, J. K., Andino, R. & Cameron, C. E. Remote site control of an active site fidelity checkpoint in a viral RNA-dependent RNA polymerase. J Biol Chem 280, 25706–25716 (2005).

27. Alanazi, A. S., James, E. & Mehellou, Y. The ProTide prodrug technology: where next? ACS Med Chem Lett 10, 2–5 (2019).

